# Recommendations for the ethical and accurate use of population descriptors: a trainee-led survey of early-career researchers

**DOI:** 10.64898/2026.07.01.735829

**Authors:** Jayati Sharma, Betzaida Maldonado, Rachel Ungar, Alvina Adimoelja, JP Flores, Tamara Gjorgjieva, Krystin Jones, Alyna Khan, Diane Xue, Roshni Patel, Christa Caggiano

## Abstract

Despite the importance of population descriptors in human genomics research, many scientists struggle to translate evolving ethical guidelines into their computational workflows. To characterize this gap between recommendations and implementation, we conducted a mixed-methods survey of early-career researchers to assess how they understand and implement the landmark 2023 NASEM report on the use of population descriptors in human genetics research. We show that while exposure to the report fosters ethical awareness, fundamental misconceptions about race and ancestry persist across academic disciplines, and trainees face structural bottlenecks, including legacy data constraints and a lack of technical confidence. To address this gap, we offer actionable, stakeholder-specific recommendations across the research lifecycle ranging from decision-support tools to “bring-your-own-data” workshops to leadership from academic journals, scientific societies, and trainee mentors. Ultimately, we argue that to promote scientific rigor and reduce bias in genetic discoveries, the scientific ecosystem must invest in the infrastructure necessary to empower the next generation of researchers.

## Introduction

The use of population descriptors in human genetics and genomics research has come under increased scrutiny in recent years. Researchers often use population descriptors such as race, ethnicity, and genetic ancestry to group research participants—either to descriptively characterize a dataset or to inform downstream analyses. These practices span a wide range of research areas, including gene discovery, prediction of disease risk, studies of human evolutionary history, and efforts to understand health disparities.^1–5^

Though certain uses of population descriptors can provide scientific value, inappropriate use can hamper scientific rigor. Use of coarse or continental-level descriptors can introduce bias, reduce research reproducibility, and potentially limit the scope of genetic discoveries. In addition, the use of race, ethnicity, and even genetic ancestry as categorical proxies for genetic variation misrepresents the complexity and continuous nature of human genetic variation, which can reinforce existing social biases and beliefs about human differences and contribute to health disparities.^6,7^

Given the scientific and ethical implications associated with using population descriptors, researchers across disciplines, from geneticists to bioethicists to sociologists, have raised alarm at their inconsistent and inaccurate use. In response, multiple groups have issued recommendations or guidelines for the ethical use of population descriptors.^8–13^ Notably, in 2023 the National Academies of Science, Engineering, and Medicine (NASEM) convened an interdisciplinary committee that issued a comprehensive set of recommendations for the use of population descriptors across a wide range of study types (referred to here as the “NASEM report”).^14^ Briefly, the NASEM report recommends researchers avoid typological thinking by limiting the use of categorical ancestry labels where possible; replacing the use of race with context-specific descriptors, such as genetic ancestry or geographic origin; and providing a clear justification for how population descriptors align with the underlying biological or social variables of interest.^14^

The NASEM report drew considerable attention from the human genetics community—owing both to the high-profile nature of the commission and its broad relevance to researchers working with human genetic and genomic data—and citations of the report have continued to climb. The following year, interest in the NASEM report resurged in the wake of the high-profile backlash to the 2024 flagship publication for the All of Us Research Program’s genetic dataset, which visually conflated genetic ancestry with race in several figures.^15^ More recently, researchers have begun using the NASEM report to create actionable data models and guidelines for different stakeholders in the scientific ecosystem.^11,16^

Despite the importance of the NASEM report, little is known about the extent to which researchers have operationalized the report’s guidelines and whether they experience any barriers to doing so. There is a particular need for understanding the attitudes and experiences of early-career researchers because relative to other career stages, early-career researchers are often directly involved in the analysis of genetic data and the preparation of manuscript results. This means they play a pivotal role in the assignment of population descriptors to research subjects and the use and interpretation of these population descriptors in downstream analyses. At the same time, early-career researchers have spent less time in the field relative to senior researchers and may have less exposure to the nuances of population descriptors and how best practices have changed over time. As members of this demographic, we also recognize that early-career researchers represent an essential and effective point for intervention: the early-career researchers of today will become the senior researchers of tomorrow. Moreover, our past work demonstrates that there are meaningful opportunities to improve early-career researcher education.^17^

To learn more about the use of population descriptors by researchers who directly work with human genetic data, we conducted a mixed-methods survey of 59 early-career researchers as an exploratory analysis to understand trends in the field. Specifically, we sought to understand: (1)how early-career researchers use population descriptors and their motivations in doing so; (2)the extent to which early-career researchers understand and implement the NASEM report in their work; and (3) the challenges these researchers face in doing so. Integrating the perspectives of the survey participants, we conclude by offering recommendations to increase uptake of the NASEM report by early-career researchers. Importantly, our recommendations are aimed not only at early-career researchers themselves, but also at principal investigators, department heads, directors of graduate education, and other institutions involved both in training early-career researchers and shaping field-wide norms.

### Survey of Early-Career Researchers

#### Survey design

An anonymous survey was designed to capture the viewpoints of early-career researchers working on human genetics research that uses population descriptors. Accordingly, we restricted the survey participants to adults who had worked on analyzing human genetic data in the last three years and who identified as undergraduates, post-baccalaureate scholars, graduate students, postdoctoral fellows/residents, and research scientists or faculty with <= 3 years in that role at academic institutions or companies. Participants who reported a career stage not within this definition, or who were under the age of 18, were excluded.

Survey questions were grouped into five sections: (1) screening, (2) views and knowledge of population descriptors, (3) perception of the NASEM report, (4) participant recommendations, and (5) demographics and research background. In total, the anonymous survey consisted of 29 questions and took approximately 15-20 minutes to complete (see Supplement for more details).

#### Participant demographics

In total, 59 responses met survey criteria. The majority of participants reported working in academia (51% PhD students; 24% postdoctoral fellows; 10% early-career faculty) (Supplementary Figure 1A-B), and a small proportion worked in non-profit or industry settings (Supplementary Figure 1B). Participants spanned various scientific subfields within genetics, with 89% stating that their primary mode of research was “dry-lab” (Supplementary Figure 1C-D). Participants also spanned a range of demographic groups: 44% self-identified as White/European, 23% as Asian/Asian-American, and 13% as Hispanic/Latino (Supplementary Figure 1E), and about half of participants self-identified as a woman (Supplementary Figure 1F). Overall, respondents represented diverse backgrounds, enabling us to characterize a variety of perspectives on population descriptor use.

#### Participant responses reveal misconceptions despite ethical intent

We were first motivated to understand how participants use population descriptors in their daily research work. We asked participants how they assigned population descriptor labels to a study, allowing them to select multiple responses to this question. Most survey participants responded that they used labels defined by the dataset, cohort, or consortia used in their analyses (n = 45, 76%) (Supplementary Figure 2). Many participants also reported that they assigned labels based on genetic similarity to a reference dataset (n = 36, 61%) and nearly half based on self-reported information (n = 29, 49%). Participants were asked to rank their top priority in choosing population descriptors. The most frequent selections were improved precision of genetic discoveries (n = 8, 14%), improved generalizability (n = 7, 12%), and ethical considerations (n = 6, 10%) (Figure 1A). We also asked participants what factors they consider when choosing population descriptors. 71% of participants agreed that researchers should consider how historical misuse of population descriptors has contributed to health inequities. A majority of participants also held the belief that researchers should clearly explain how population descriptors were used in a particular study (Figure 1B) and that researchers should strive for accurate and explicit language to avoid harmful misconceptions (Figure 1C).

**Figure 1:**
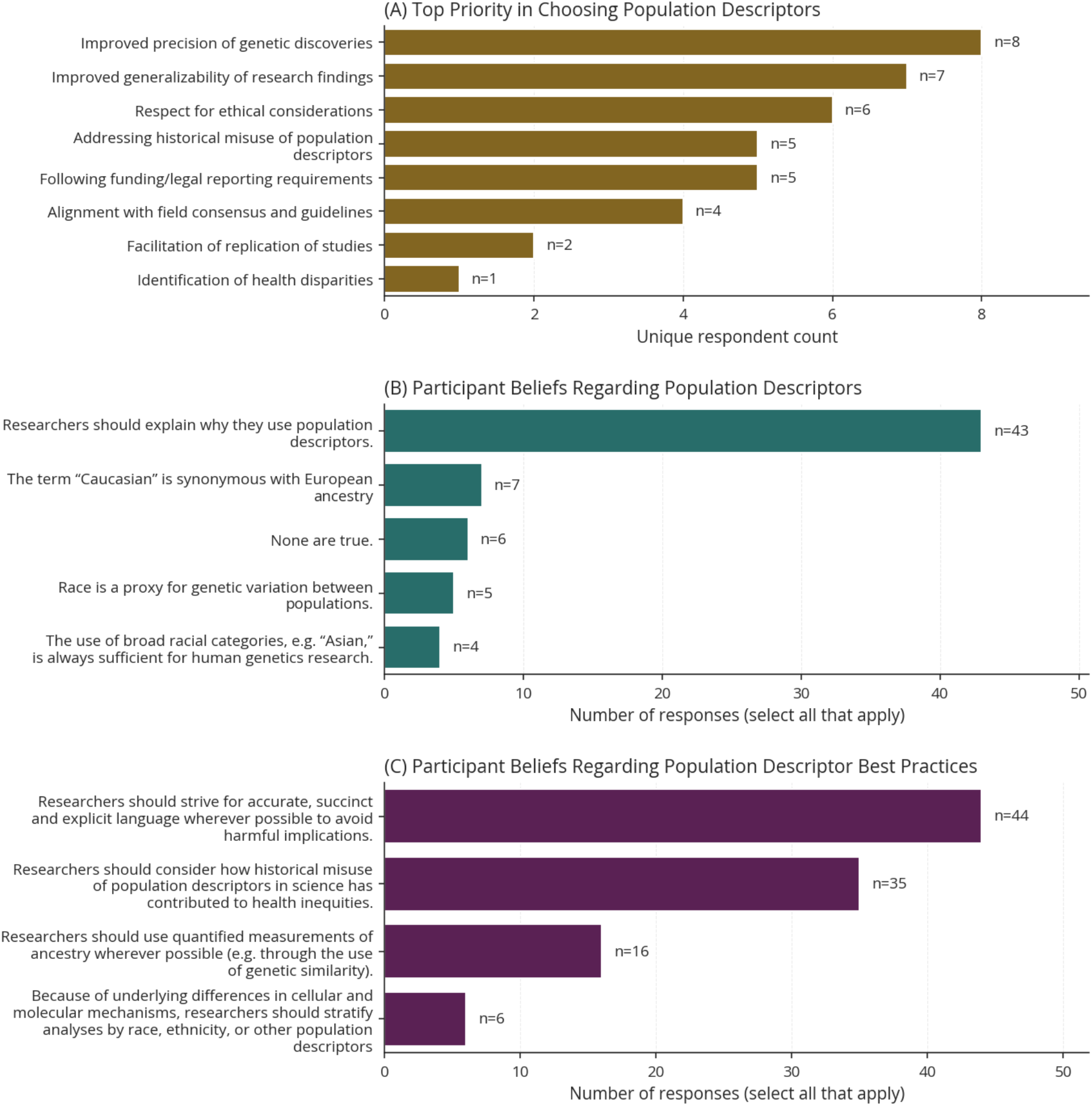
Participants prioritized precision and generalizability in choosing population descriptors, though beliefs about race as a genetic proxy and the sufficiency of broad racial categories demonstrate lingering misconceptions. (A) The top-ranked priority of participants when choosing population descriptors in their research. Participants selected one priority. (B) Participant beliefs regarding how population descriptors might be used. Participants could select multiple options. (C) Participant beliefs around the ethical considerations and best practices in population descriptor use. Participants could select multiple options.

To characterize potential biases and misconceptions, we next wanted to understand participants’ knowledge of the use of race, ethnicity, and genetic ancestry as population descriptors. We identified several common and harmful misconceptions highlighted by the NASEM report and asked participants to select which they believed to be true (“select all that apply”). Asked across multiple questions, these misconceptions included the incorrect statements that race is a proxy for genetic variation (n = 7, 12%), that “Caucasian” is synonymous with European ancestry (n = 5, 8%), that broad racial categories are always sufficient for research (n = 4, 7%), and that biological differences between groups motivate the use of race and other population descriptors (n = 6, 10%). Eleven (19%) distinct individuals expressed agreement with at least one misconception.

#### NASEM exposure fosters ethical awareness, but opportunities exist to improve implementation

Characterizing whether and how early-career researchers use the NASEM report in their work is an important step in developing frameworks for its use. Ten participants (17%) reported reading the report in its entirety, and an additional 30 participants (51%) reported that they had attended presentations on the report, read it in parts, or had generally heard of it (Supplementary Figure 3). Given that even informal exposure to the NASEM report may influence researchers’ use of population descriptors, we analyzed both subgroups together, for a total of 40 NASEM-exposed participants (64%) and 19 NASEM-unexposed participants (32%).

We found that there were slight differences in NASEM exposure across demographic/career groups (Supplementary Figure 4). For example, 70% of people identifying as a woman reported NASEM exposure versus 52% of those identifying as a man (Supplementary Figure 4B).

Approximately 70% of PhD and postdoctoral fellows had exposure to the report, compared to only 33% of early-career faculty (Supplementary Figure 4C). Furthermore, there were apparent differences by field of study. No one who selected computer science as their field of study (n = 3) was familiar with the report (Supplementary Figure 4E). However, these differences should be understood in the context of small sample sizes.

Next, we assessed the impact of the NASEM report on participants’ usage of population descriptors, knowledge of population descriptors, and understanding of best practices. Only 15 of 40 NASEM-exposed individuals (38%) reported using the report’s guidelines to assign population descriptors (Supplementary Figure 2). Moreover, we found that fundamental misconceptions about race, ethnicity, and ancestry persisted regardless of exposure to the NASEM report. However, NASEM-exposed participants believed misconceptions slightly less frequently than NASEM-unexposed participants. For example, 4 (21%) unexposed participants incorrectly believed “Caucasian” is synonymous with European ancestry compared to 3 (8%) exposed participants (Supplementary Figure 5).

Interestingly, we observed that exposure to the NASEM report had an association with researchers’ perception of the ethical implications of using population descriptors. Only NASEM-exposed individuals ranked “respect for ethical considerations” as a top priority regarding the use of population descriptors (Supplementary Figure 6). We observed similar trends when we asked participants to self-report their understanding of the potential benefits and harms of using population descriptors. NASEM-exposed participants were more likely to report that they understood these ethical implications “well” or “very well” (Supplementary Figure 7).

#### Trainees face barriers in implementing the NASEM guidelines

We next sought to more clearly define the NASEM report’s utility to participants and to identify potential barriers to implementation. While the majority of respondents (55%) found the report moderately to extremely useful, and 69% reported a strong personal understanding of its contents (Supplementary Figure 8A-B), this perceived utility did not translate into their ability to implement the report’s guidelines. In fact, only 29% of participants reported high or very high confidence in their ability to apply the recommendations to their own research (Supplementary Figure 8C). Open-ended survey responses suggest this gap is driven by a lack of confidence in applying guidelines; concerns over co-author and reviewer scrutiny; and the technical limitations of legacy datasets that lack the granularity required for population descriptor reform. For example, one respondent shared, “I often get comments from coauthors confused when using ‘new’ labels and asking if they correspond to the ‘old’ labels they are accustomed to.”

A majority of respondents (n = 35, 59%) indicated that a step-by-step technical guide for applying NASEM recommendations to real-world data would be the most effective motivator for implementation, followed closely by requirements by journals and funding agencies (Figure 2A). Even among NASEM-exposed participants, 23 participants (58%) desired a technical guide for implementing the recommendations (Supplementary Figure 9). This need for concrete, actionable resources directly mirrors the primary challenges reported among all participants: population descriptors being pre-decided (n = 18), a lack of confidence in applying the guidelines (n = 14), and data not having sufficient information to implement the guidelines (n = 13) (Figure 2B).

**Figure 2:**
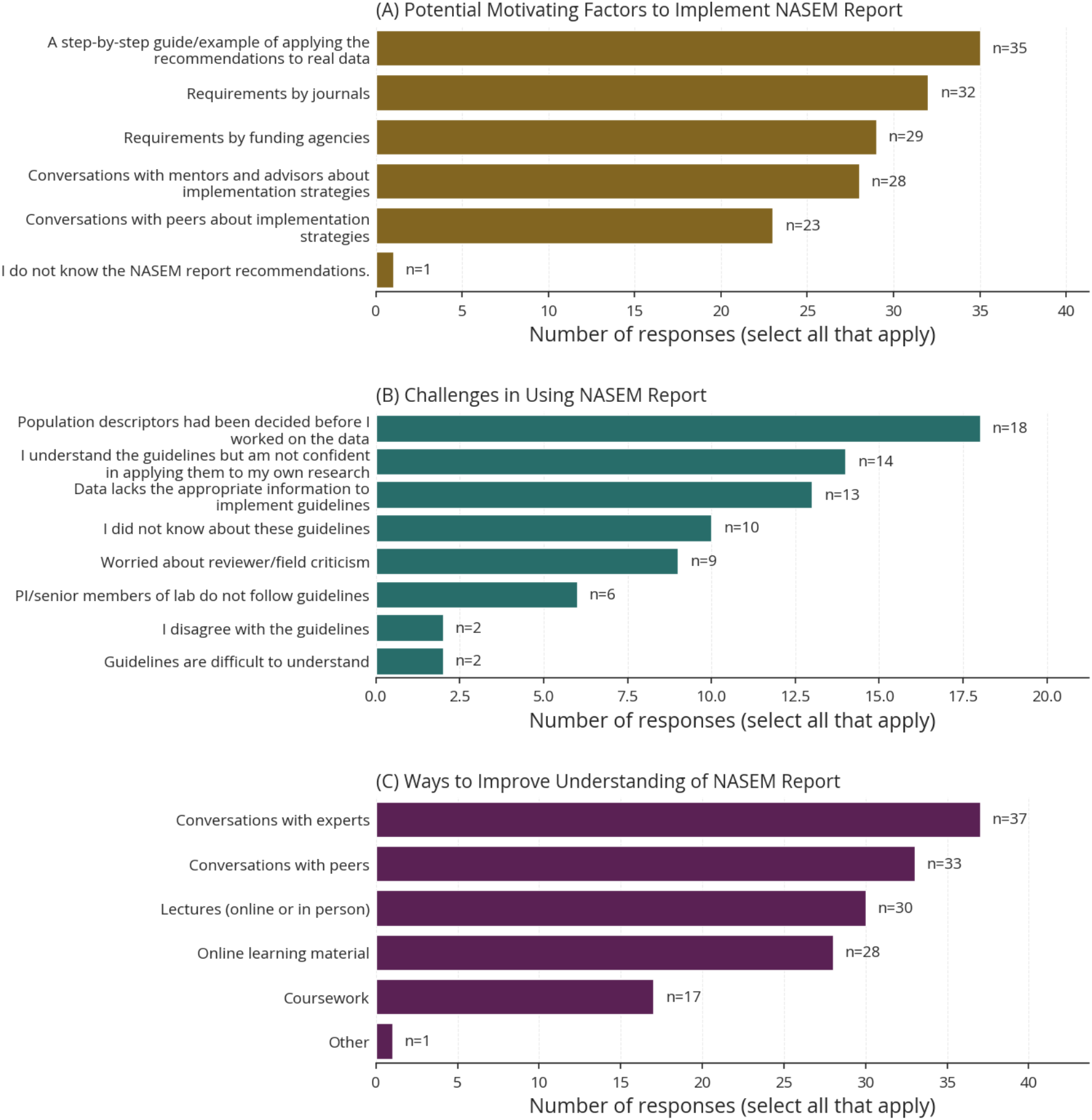
Despite broad awareness of the NASEM report, participants report structural barriers and a lack of confidence in implementation and prefer interpersonal and expert-driven support to bridge that gap. (A) Factors that would motivate participants to implement the NASEM report recommendations. (B) Challenges participants identified when attempting to implement the guidelines in their own research. (C) Ways that participants would like to improve their understanding of the NASEM report recommendations. Participants could select multiple responses for all questions.

#### Ethics training accelerates literacy but fails to prevent misconceptions

To understand possible areas where training gaps exist, we categorized participants into those who had and not been formally trained in ethics broadly. We defined “formal” ethics training as respondents who had received any formal training in ethics or bioethics: trainings in the Ethical, Legal, and Social Implications of research (ELSI), Science & Society or Science & Technology Studies (STS), or the Controlled Access Tier Training from the All of Us Research Program (n = 46, 78%). Mandatory Responsible Conduct of Research (RCR) classes were not included in our definition given these classes are extremely common in early-career training and do not focus on human genetic data. As expected, we observed that a high proportion of our sample had formal ethics training across fields and other demographic categories (Supplementary Figure 10).

Participants who had formal ethics training were more likely to correctly respond to objective questions regarding population descriptors (Supplementary Figure 11). However, misconceptions persisted regardless of training. We also observed field-specific trends, with respondents from statistics or computer science backgrounds more likely to have misconceptions (n = 12, 46%) than human genetics/bioinformatics (n = 13, 31%) or public health/bioethics (n = 4, 36%), potentially reflecting divergent disciplinary norms (Supplementary Figure 12). While formal training improves alignment with NASEM guidelines, these data demonstrate that conceptual gaps persist across all research backgrounds, and early-career researchers across disciplines could benefit from additional training.

### Recommendations

Based on the perspectives gathered in our survey, we identified a persistent gap between researchers’ ethical intent and research practice with respect to population descriptors. Despite broad awareness of the NASEM report and genuine motivation to apply its recommendations, survey participants demonstrated lingering misconceptions about race, ethnicity, and genetic ancestry and reported low confidence in implementing guidelines. They also identified structural barriers, including legacy data constraints and insufficient institutional support, that impede their implementation of the NASEM guidelines.

This pattern echoes findings from the analogous domain of sex and gender reporting in biomedical research, where existing work has demonstrably shifted researcher attitudes without producing commensurate changes in publication practice.^18–21^ Our results suggest that the same risk applies to population descriptor reform: without mechanisms that embed guidelines into the daily workflows of researchers, normative shifts in attitude may not yield the methodological changes the field requires. Taken together, our findings suggest that the field must invest in concrete infrastructure to bridge the implementation gap.

To address this need, we leveraged suggestions from survey participants and our own experience as early-career researchers to identify key stakeholder-specific recommendations at multiple stages of the research lifecycle. In some places, our recommendations reiterate those made by existing publications or workshops, including the 2023 NASEM report studied here, the 2024 NASEM report on “The Use of Race and Ethnicity in Biomedical Research,” and the 2024 National Human Genome Research Institute (NHGRI) workshop, “Population Descriptors for Legacy Genomic Data: Challenges and Future Directions.”^22^ In addition, we include several original recommendations denoted as such (Table 1). We discuss five broad themes of our recommendations below.

**Table 1.**
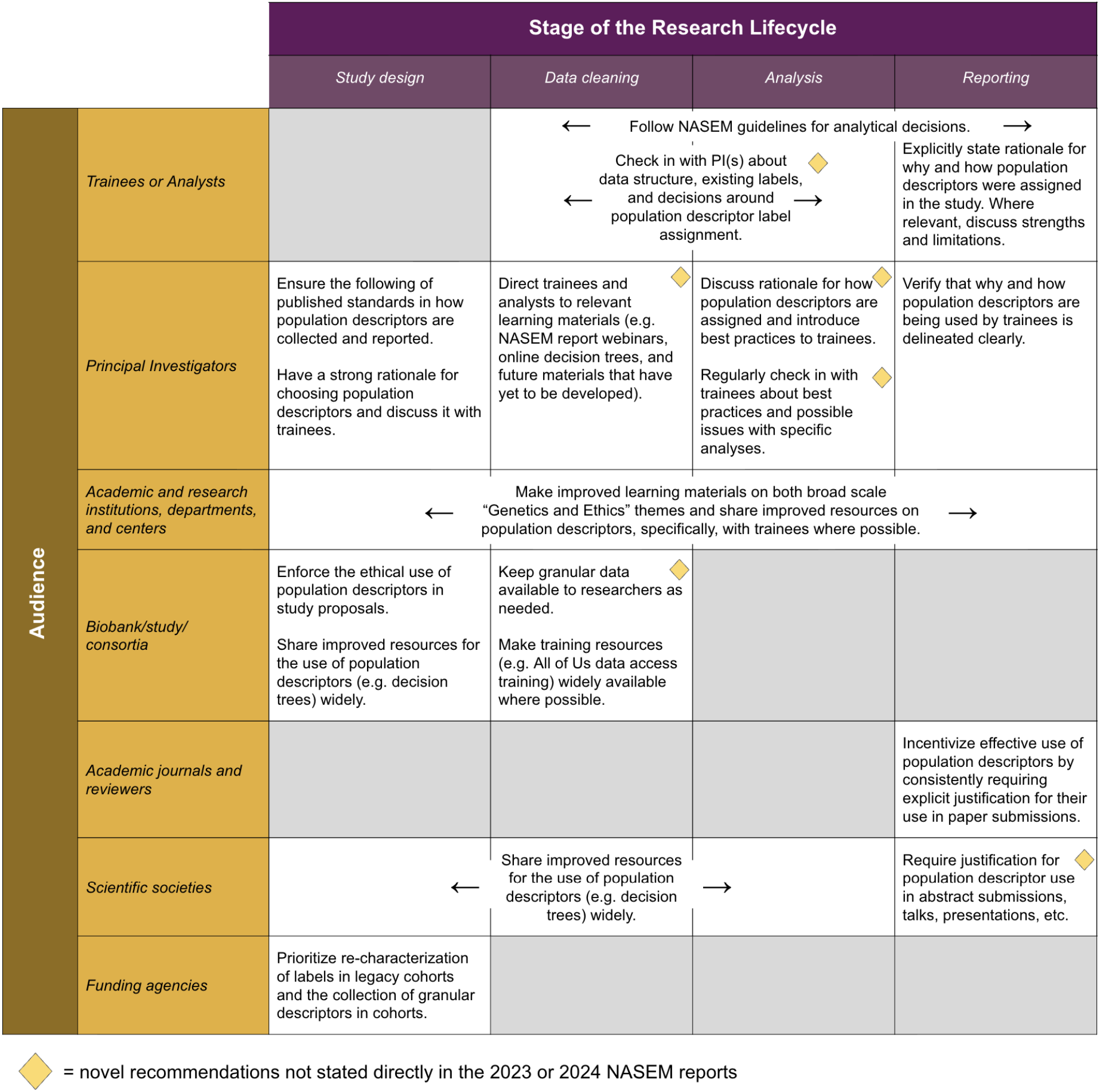
Recommendations for the effective use of population descriptors.

First, we recommend the development of structural incentives to encourage the accurate and ethical usage of population descriptors. Some of participants’ reported reliance on outdated labels in source data stems from analytical inertia rather than technical limitations. At times, survey participants reported not feeling empowered to determine data labels on their own. Other participants cited field consensus and reviewer criticism as important concerns in their decisions to implement NASEM guidelines or not. Reiterating the recommendations of Feero et al., academic journals publishing computational human genetic research should require justification for the use of population descriptors in manuscript submissions.^23^ Scientific societies should encourage the same in abstract and oral presentations. Earlier in the research lifecycle, consortia and biobanks should likewise require clear justification in data use applications. Such requirements directly shape how trainees learn to communicate science: when manuscripts must include explicit justification for population descriptor choices, trainees engaging in the writing process are prompted to think critically about these decisions at a formative stage.

Second, our survey identified a legacy data bottleneck where many participants felt restricted by the existing and often outdated features of their source data. Trainee education should enable them to analyze study and questionnaire design to understand what population descriptor data exist, how they were collected, and limitations therein. Where possible, trainees and PIs can discuss ways to exercise analytical agency by deriving more precise descent-associated descriptors, such as through fine-scale population structure analysis or local ancestry inference, or by using modern statistical methods that avoid discrete ancestry categories.^24–26^ When such analysis is not technically feasible or appropriate, researchers should explicitly acknowledge the limitations of legacy descriptors, including how these broad categories may obscure internal diversity or introduce confounding. While trainees and PIs can effect such changes at an individual level, broader institutional action is also necessary. Accordingly, biobanks or consortia should make granular, multi-level population descriptors available for existing datasets and in the interim, should provide clear guidance and training on the use of existing descriptors.

Funding agencies should also prioritize the recharacterization of legacy cohorts to harmonize descriptors across studies. These efforts should be grounded in community perspectives to avoid prescribing labels without sufficient understanding of the populations they represent.

Our final three recommendations span different aspects of trainee education. Respondents explicitly called for expert-led conversations and guided learning, which suggests that the current self-study model is insufficient for mastering these complex definitions. Instead, leadership is needed from the individuals and institutions responsible for training early-career researchers. Thus, we specifically recommend targeted resources for use in trainee research, leadership from trainee mentors, and changes to trainee coursework.

Given that a majority of participants voiced a desire for a step-by-step guide on population descriptors, the field clearly needs targeted resources that can be integrated into trainees’ day-to-day research. One example of such a resource is the All of Us Research Program’s training module on ethical considerations, which provides guidance on applying population descriptors in this dataset and is required of researchers working with data from diverse populations. In addition, there is a need for concrete decision-support tools that expand beyond the decision trees provided by the NASEM report. For example, a centralized toolkit with open-source code and standardized reporting templates could increase transparency and reproducibility around researcher-derived population descriptors and provide trainees with a targeted resource to aid in implementing the guidelines in their workflows. Recent publications have begun to provide such examples, but scientific societies and academic institutions should prioritize the further development and dissemination of such materials.^11,16^ Finally, we recommend “bring-your-own-data” workshops at conferences where researchers can gain the practical knowledge and confidence to apply NASEM guidelines to their own datasets in a sandbox environment.

Fourth, principal investigators (PIs) and other trainee mentors should initiate regular discussions about the use of population descriptors in trainee projects. PIs should also engage their labs in trainings on the ethical and accurate use of population descriptors. This can happen at minimal cost and provide profound benefits: PIs play a critical role in shaping trainee confidence. By initiating open and early conversations about population descriptor choices at project initiation, mentors can normalize these decisions as a standard part of rigorous research practice rather than an afterthought.

Fifth, department heads and directors of graduate education should prioritize the development of interdisciplinary “Genetics and Society” curricula that move beyond standard research ethics training. Courses on the broader social and ethical implications of human genetics have already been successfully implemented in the Stanford Department of Genetics and the UNC Biological and Biomedical Sciences Program for bioscience PhD students.^17^ Future course development should also enable trainees to engage in practical exercises, such as deconstructing legacy labels like “Caucasian” and replacing them with geographically or genetically precise alternatives.

## Discussion

Here we present findings from a survey of 59 early-career researchers on their use of population descriptors and the impact of a landmark report commissioned by the National Academies. As the individuals often directly responsible for implementing population descriptor assignments in day-to-day analyses—many of whom will serve as future leaders in the field—early-career researchers occupy a critical leverage point for reform. Yet, our survey documents a group that feels simultaneously motivated by the NASEM report and constrained by their context: by legacy data features they did not choose, by datasets without sufficient information to implement NASEM recommendations, by training programs that do not translate broad ethics education into methodological guidance, and by peer pressures that discourage deviation from field conventions.

Overall, we find that while self-reported exposure to the NASEM report is associated with increased perception of the importance of ethical and accurate population descriptor usage, this may not necessarily translate into action. This discrepancy suggests that early-career researchers may lack the technical or practical ability to apply theoretical concepts, highlighting the need for targeted pedagogical interventions to align research behavior with ethical priorities. To that end, we put forth recommendations targeted at stakeholders across the research process, ranging from consortia to principal investigators (PIs) advising trainees. Though best practices regarding population descriptors are dynamic and learning materials will inevitably require adaptation over time, developing such resources will establish a strong foundation for future iteration.

The ethical and accurate use of population descriptors is fundamentally intertwined with conversations about equity and diversity in research. Accordingly, implementing our recommendations may seem to require sustained investments in infrastructure that currently faces significant political headwinds. Yet, the scientific argument for these investments can be made without invoking diversity or equity: conflating race with genetic variation and defaulting to coarse population labels introduces bias, reduces reproducibility, and limits genetic discovery. Concretely, the resources required to implement many of our recommendations are tractable even amid opposition from federal, state, or university leadership. For example, centralized open-source toolkits and reporting templates require one-time development and can be updated by existing consortia or scientific societies’ working groups. Workshops can be embedded in existing conference programming. Population descriptor-specific training modules can be integrated into mandated researcher ethics training given buy-in from instructors.

Fundamentally, broader and more accurate population representation results in better-powered studies that enable reproducibility in other contexts, underscoring that researchers need not treat equity and scientific rigor as competing or independent aims.

Several limitations of our survey warrant acknowledgment. Our measure of NASEM exposure was self-reported and necessarily coarse, collapsing diverse forms of engagement from full reading to passive secondhand exposure into a single variable, which may have attenuated our ability to detect exposure-related differences. Our survey’s rating questions relied on a Likert scale, which, while standard for attitudinal surveys, is inherently subjective and subject to social desirability bias. Respondents may have reported stronger agreement with ethical best practices than they believe. Finally, our sample of 59 respondents is small and self-selecting: researchers who chose to complete a survey on population descriptors likely differ systematically from those who did not, potentially inflating estimates of awareness and ethical concern. At the same time, identifying misconceptions among these participants highlights the pervasiveness of this issue: there is room for growth even in individuals with high preexisting motivation and/or training.

Owing to our small sample sizes, our ability to understand discipline-specific trends was limited. Though we observed some differences—for example, participants with computer science or statistics backgrounds were more likely to hold incorrect beliefs—further work is needed to develop larger, diverse cohorts to fully characterize such trends in a larger study population.

Regardless, our data reiterate the importance of interdisciplinary educational resources: misconceptions about race and ancestry persisted regardless of academic background.

Finally, a crucial area for research is to evaluate how researchers’ attitudes and practices evolve over time. The NASEM report is only a few years old, and human genetics research has a long history. In the future, longitudinal surveys and bibliometric methods will be able to characterize the impact of the NASEM report more comprehensively. Future work should also aim to characterize the attitudes and practices of senior researchers. Interventional studies that examine how the recommendations suggested here impact the beliefs and practices of early-career researchers over time will also be helpful in developing future guidelines.

Ultimately, our findings highlight a critical pivot point: the scientific ecosystem must move from passively disseminating guidelines to actively building the technical infrastructure necessary for their adoption—namely, by developing computational toolkits, mandating transparent reporting, and iteratively evaluating researcher attitudes and resource development. Together, these investments can empower early-career researchers to make analytical choices that align with field ethics, ensuring genomic analyses accurately reflect the complexity and continuous nature of human genetic variation. The transition toward the ethical and accurate use of population descriptors is a structural necessity for human genomics.

## Ethical Considerations

Before administering the survey, we received approval from the Mount Sinai Health System IRB (protocol STUDY-24-01473). No personal identifying information was gathered through the fully anonymized survey. The survey was not distributed directly to potential respondents by authors to limit the authors’ knowledge of participants’ identities and to prevent biasing of the sample of respondents. The survey remained open for 10 weeks, and the resulting data was only directly accessed by one author for data cleaning.

## Acknowledgements

The authors thank all survey participants for their time and thorough responses. We also thank the American Society of Human Genetics, NHGRI T32 training programs, administrative program staff, and the scientific communities on social media platforms who supported survey dissemination. J.S. was supported by 1F31HG013440. R.U. was supported by a T32 grant 5T32HG00895310. C.C. was supported by F32HG014400. We would also like to thank Eimear Kenny, Daphne Martschenko, and Genevieve Wojcik for their thoughtful comments on a draft version of this piece.

## Author contributions

Project and team administration - J.S.; survey drafting - J.S., R.P., A.A., R.U., and C.C; survey piloting - D.X., B.M., and T.G.; survey dissemination - all authors; IRB protocol development - C.C., B.M., and R.U.; data storage & formal analysis - C.C.; writing (original draft) - J.S., C.C., R.P., and B.M.; writing (reviewing and editing) - all authors.

## Data availability

Our raw, unidentifiable survey data and code used for analysis are available at DOI: 10.5281/zenodo.21045270

