## Supplement for "Recommendations for the ethical and accurate use of population descriptors: a trainee-led survey of early-career researchers"

### Supplemental Note

#### Participant Demographics

For the demographic questions, race, ethnicity, and gender identity labels were generally aligned with the 2024 NIH Policy on Collection of Race and Ethnicity Data's revised demographic collection standards and the American Community Survey's gender identity questions. We collected this self-reported information to understand the demographic composition of participants, which resembled the broader human genetics field

For reference, see: [https://www.ashg.org/wp-content/uploads/2022/11/WorkforceSurveyReport_Report_FINAL2.pdf](https://www.ashg.org/wp-content/uploads/2022/11/WorkforceSurveyReport_Report_FINAL2.pdf).

#### Participant Recruitment

The survey was widely distributed through scientific networks frequented by early-career scientists, including the American Society of Human Genetics, National Human Genome Research Institute T32 institutions, institutional/department email listservs, Slack channels, and social media (X, Bluesky, LinkedIn, Facebook, and Mastodon). To reduce the chance of coercion, no individual messages were sent to any specific scientist, and a maximum of three emails or Slack posts were sent to each group. Recruitment was open for 10 weeks during the summer of 2025.

#### Survey Design

The survey was designed by J.S., R.P., A.A., R.U. and piloted by D.X., B.M., and T.G. to refine questions, length, and content.

#### Survey Questions

##### 1. SCREENING

The purpose of the following few questions is to determine your eligibility for this study.

**Are you 18 years or older?**

1. Yes
2. No

(if answer is NO)

*Thank you for your interest in this survey. Based on your responses, you do not meet the eligibility criteria at this time. This concludes your participation, and we appreciate your time.*

2. **What career stage are you currently in?**

1. Undergraduate
2. Post-baccalaureate
3. Graduate (Master's)
4. Graduate (PhD)
5. Graduate (MD)
6. Post-graduate (Post-doctoral fellow)
7. Post-graduate (Resident)
8. Tech/scientist (academic) < 3 years in this role
9. Tech/scientist (academic) ≥ 3 years in this role
10. Tech/scientist (industry) < 3 years in this role
11. Tech/scientist (industry) ≥ 3 years in this role
12. Faculty < 3 years in this role
13. Faculty ≥ 3 years in this role
14. Other

(if answer is I, K, M )

*Thank you for your interest in this survey. Based on your responses, you do not meet the eligibility criteria at this time. This concludes your participation, and we appreciate your time.*

3. **Do you currently work with or have you recently worked with (in the last 3 years) the computational analysis of human genetics/genomics data at the population level?**
    1. Yes
    2. No

(if answer is NO)

*Thank you for your interest in this survey. Based on your responses, you do not meet the eligibility criteria at this time. This concludes your participation, and we appreciate your time.*

---

Throughout the survey, we will refer to this term "population descriptors" regularly. For reference, a population descriptor is "a concept or classification scheme that categorizes people into groups (or "populations") according to a perceived characteristic or dimension of interest." Population descriptors can include, but are not limited to, labels such as race, ethnicity, and geographic location.

#### 2. VIEWS AND KNOWLEDGE OF POPULATION DESCRIPTORS

1. **How do you assign population descriptor labels to samples in your study? (Select all that apply)**
    1. Existing literature search
    2. Labels defined by the dataset/cohort/consortia under study

    3. Labels based on self-reported information
    4. Labels based on genetic similarity to a reference dataset/population
    5. Based on individual experience/best judgement
    6. My PI/senior member of lab decides this
    7. Using guidelines from the NASEM report: "*Using Population Descriptors in Genetics and Genomics Research*"
    8. I do not think about this
    9. Other: _____
2. **Which of the following do you consider the most when choosing population descriptors in your research? (Rank in order of importance).**
    1. Improved generalizability of research findings
    2. Improved precision of genetic discoveries
    3. Identification of health disparities
    4. Respect for ethical considerations
    5. Facilitation of replication of studies
    6. Alignment with field consensus and guidelines
    7. Addressing historical misuse of population descriptors
    8. Following funding/legal reporting requirements
    9. None of the above
    10. Other: _____
3. **How well do you understand the potential <u>benefits</u> of using accurate population descriptors in your research?**
    1. 1: Not at all
    2. 2: Slightly
    3. 3: Somewhat
    4. 4: Well
    5. 5: Very well
4. **How well do you understand the potential <u>harms</u> of using inaccurate or incomplete population descriptors in your research?**
    1. 1: Not at all
    2. 2: Slightly
    3. 3: Somewhat
    4. 4: Well
    5. 5: Very well

---

*This section of the survey seeks to assess your understanding of race, ethnicity, ancestry, and population descriptors in human genetics research.*

5. **Have you heard of the NASEM report titled "Using Population Descriptors in Genetics and Genomics Research"? (Select all that apply)**
    1. Yes, I have heard of it
    2. Yes, I have skimmed the report
    3. Yes, I have read this report in parts

4. Yes, I have read the report in full
5. Yes, and I have attended a presentation explaining the report
6. No
7. Not sure

(if answer is NO, complete this section and then skip to Section 4: Recommendations)

6. **Which of the following do you believe to be true? (Select all that apply)**
    1. Race is a proxy for genetic variation between populations.
    2. The term “Caucasian” is synonymous with European ancestry
    3. Researchers should explain why they use population descriptors.
    4. The use of broad racial categories, e.g. “Asian,” is always sufficient for human genetics research.
    5. None of the above are true.
7. **If you are conducting a study to characterize health disparities due to both genetic and environmental effects in a diverse US population, which kinds of population descriptors may be relevant in describing the results of your study? (Select all that apply)**
    1. Race and ethnicity labels
    2. Labels based on genetic similarity to reference populations
    3. Labels based on self-reported information
    4. Labels defined by the dataset/cohort/consortia under study
    5. None of the above
8. **Which statement <u>best summarizes</u> the ethical considerations regarding the use of population descriptors in human genetics research?**
    1. Researchers should use quantified measurements of ancestry wherever possible (e.g. through the use of genetic similarity).
    2. Researchers should consider how historical misuse of population descriptors in science has contributed to health inequities.
9. **Which statement <u>best summarizes</u> the best practices for using ancestry-related population descriptors in genetic research?**
    1. Because of underlying differences in cellular and molecular mechanisms, researchers should stratify analyses by race, ethnicity, or other population descriptors
    2. Researchers should strive for accurate, succinct and explicit language wherever possible to avoid harmful implications.

---

#### 3. SUBJECTIVE PERCEPTION OF NASEM REPORT

1. **On a scale from 1-5, how <u>useful</u> do you think the NASEM report on population descriptors is to your research**?
    1. 1: Not at all useful
    2. 2: Slightly useful
    3. 3: Moderately useful
    4. 4: Very useful

    5. 5: Extremely useful
    6. N/A: The NASEM report is not useful to my research
2. **On a scale from 1-5, how well did you understand the recommendations in the NASEM report on population descriptors?**
    1. 1: Not at all
    2. 2: Slightly well
    3. 3: Moderately well
    4. 4: Very well
    5. 5: Extremely well
    6. N/A: I did not read the NASEM report.
3. **On a scale from 1-5, how confident are you in your ability to apply the NASEM population descriptor recommendations in your research?**
    1. 1: Not at all confident
    2. 2: Slightly confident
    3. 3: Moderately confident
    4. 4: Very confident
    5. 5: Extremely confident
    6. N/A: I did not read the NASEM report.
4. **Which of the following tools from the NASEM report do you find yourself using most often? (Select all that apply)**
    1. Interactive decision tree
    2. Chapter 5 text on choosing population descriptors
    3. Published examples of genetic similarity descriptions ("1KG-AFR-like")
    4. Other: _____
    5. None of the above
5. **What challenges do you experience in assigning population descriptor labels using the NASEM guidelines in your research? (Select all that apply)**
    1. I did not know about these guidelines
    2. Guidelines are difficult to understand
    3. I understand the guidelines, but am not confident in applying them to my own research
    4. PI/senior members of lab do not follow guidelines
    5. Data lacks the appropriate information to implement guidelines
    6. Worried about reviewer/field criticism
    7. Population descriptors had been decided before I worked on the data
    8. I disagree with the guidelines
    9. I do not find assigning population descriptors challenging
    10. Other: _____
    11. N/A

---

4. RECOMMENDATIONS

1. **What would improve your (or others) understanding of the use of population descriptors in human genetics research? (Select all that apply)**
    1. Coursework
    2. Online learning material
    3. Lectures (online or in person)
    4. Conversations with peers
    5. Conversations with experts
2. **Which of the following would motivate you to implement the NASEM report recommendations in your research? (Select all that apply)**
    1. A step-by-step guide/example of applying the recommendations to real data
    2. Conversations with peers about implementation strategies
    3. Conversations with mentors and advisors about implementation strategies
    4. Requirements by funding agencies
    5. Requirements by journals
3. **Are there any other recommendations you have for improving your and others' ability to implement the NASEM report's recommendations?**
    1. Free form response: _____

#### 5. DEMOGRAPHICS & RESEARCH BACKGROUND

1. **Gender and gender identity (select all that apply)**
    1. Gender non-binary: Gender nonconforming, genderqueer, nonbinary
    2. Man
    3. Woman
    4. Transgender or trans
    5. Other: _____
    6. Prefer not to respond
2. **Race and/or Ethnicity (select all that apply)**
    1. American Indian or Alaska Native (e.g., Navajo Nation, Blackfeet Tribe, Inupiat Traditional Gov't., etc.)
    2. Asian or Asian American (e.g., Chinese, Japanese, Filipino, Korean, South Asian, Vietnamese, etc.)
    3. Black or African American (e.g., Jamaican, Nigerian, Haitian, Ethiopian, etc.)
    4. Hispanic or Latino/a (e.g., Puerto Rican, Mexican, Cuban, Salvadoran, Colombian, etc.)
    5. Middle Eastern or North African (e.g., Lebanese, Iranian, Egyptian, Moroccan, Israeli, Palestinian, etc.)
    6. Native Hawai`ian or Pacific Islander (e.g., Samoan, Guamanian, Chamorro, Tongan, etc.)
    7. White or European (e.g., German, Irish, English, Italian, Polish, French, etc.)
    8. My race or ethnicity is best described as: _______ (Feel free to use the text box and/or you can simply select categories above.)
    9. Prefer not to respond
3. **Type of Institution (select all that apply)**
    1. Research university

    2. Medical school or hospital
    3. Liberal arts college or primary undergraduate institution
    4. Government
    5. Non-profit
    6. Industry
    7. Other: _____
4. **This survey will cover topics from a report published by a US organization. Is your institution, organization, or company based in the US?**
    1. Yes
    2. No
5. **Primary mode of research (Select all that apply):**
    1. Computational/"dry"
    2. Experimental/"wet"
6. **Field of Study (Select all that apply):**
    1. Molecular Biology and/or Human Genetics
    2. Computer Science
    3. Bioinformatics
    4. Epidemiology and/or Public Health
    5. Statistics, Biostatistics, and/or Data Science
    6. Anthropology, Social Science Medicine, Law, and/or Bioethics
    7. Clinical or Medical Genetics
7. **Primary Scientific Interest (from ASHG) (Select one):**
    1. Cancer
    2. Complex traits
    3. Developmental genetics
    4. Epigenetics
    5. Evolutionary and population genetics
    6. Genetic counseling, ELSI
    7. Genetic therapies
    8. Genetic resources and databases
    9. Mendelian traits and rare diseases
    10. Molecular effect of genetic variation
    11. Statistical genetics
8. **Does the genetic data you work with contain individuals across racial, ethnic, or genetic ancestry groups?**
    1. Yes
    2. No
9. **Have you received training in any of the following topics? (via courses, workshops, seminars, etc) (Select all that apply):**
    1. Ethics or Bioethics
    2. Ethical, Legal, and Social Implications (ELSI)
    3. NIH requirement Responsible Conduct of Research (RCR)
    4. Science & Society or Science & Technology Studies (STS)
    5. All of Us Controlled Access Tier Training

6. Informal training [conversations with advisors/mentors; discussion in lab/dept meetings; social media explainers]
7. None of the above

### Supplemental Figures

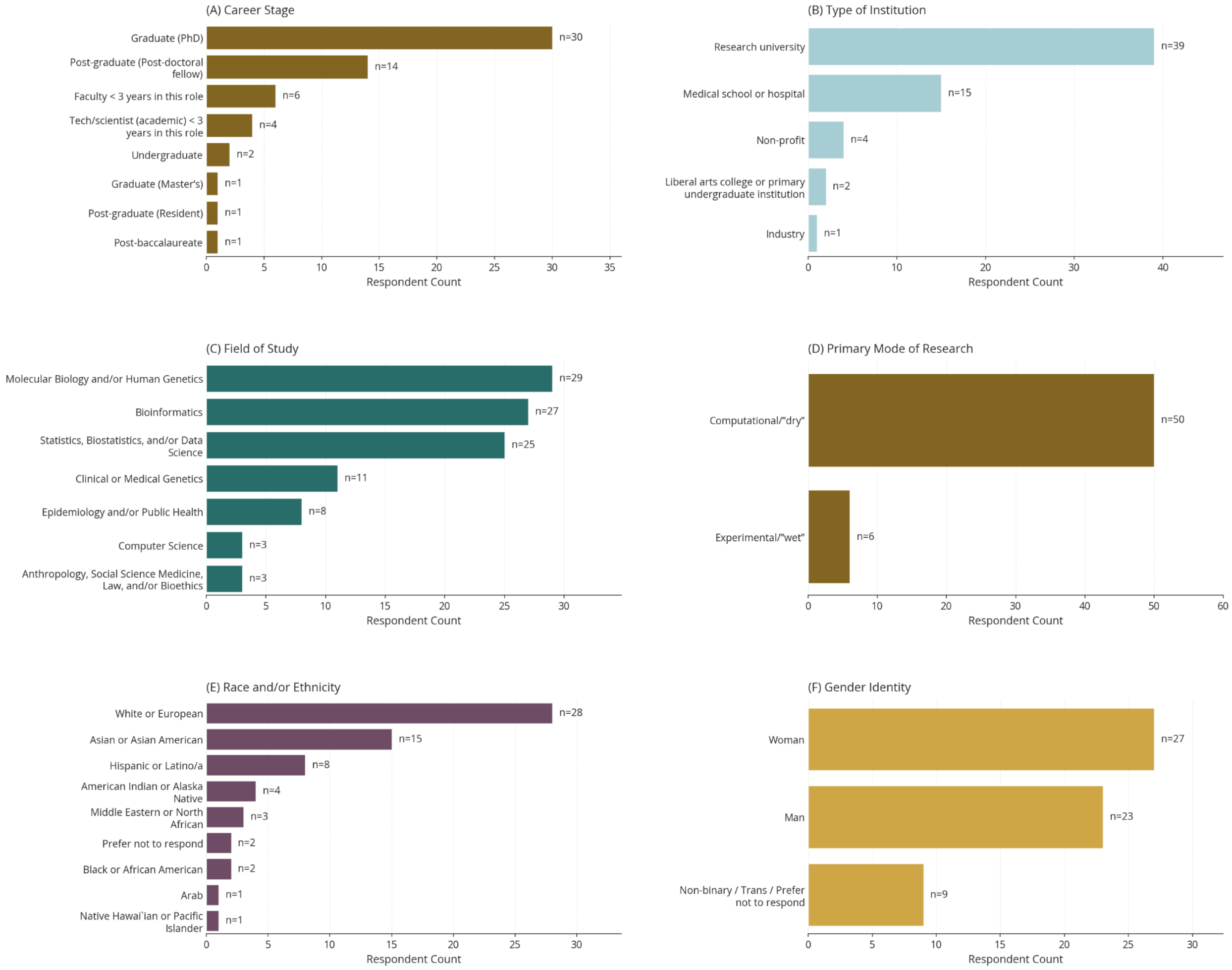


**Supplementary Figure 1:** *Participant demographics.* For each participant that chose to respond to demographic questions, the count of those (A) at each career stage, (B) at type of institution, (C) in a specific scientific field, (D) that perform computational or wet lab research; and the participant (E) self-reported race/ethnicity and (F) gender identity. Participants were allowed to select multiple responses when they felt it was appropriate.

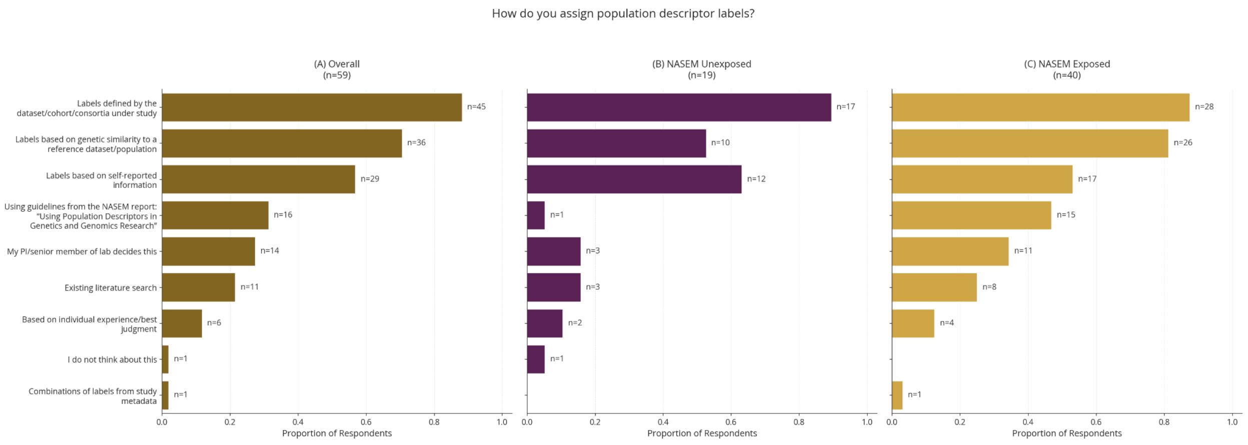


**Supplementary Figure 2:** *How participants assign population descriptors.* The proportion of participants selecting each answer to the question, *"How do you assign population descriptor labels?"* for (A) the entire cohort, (B) participants reporting that they were exposed to the NASEM report, and (C) participants reporting they were not exposed to the NASEM report. Participants were allowed to select multiple answers; thus, the total proportions in a given category can be greater than 1.

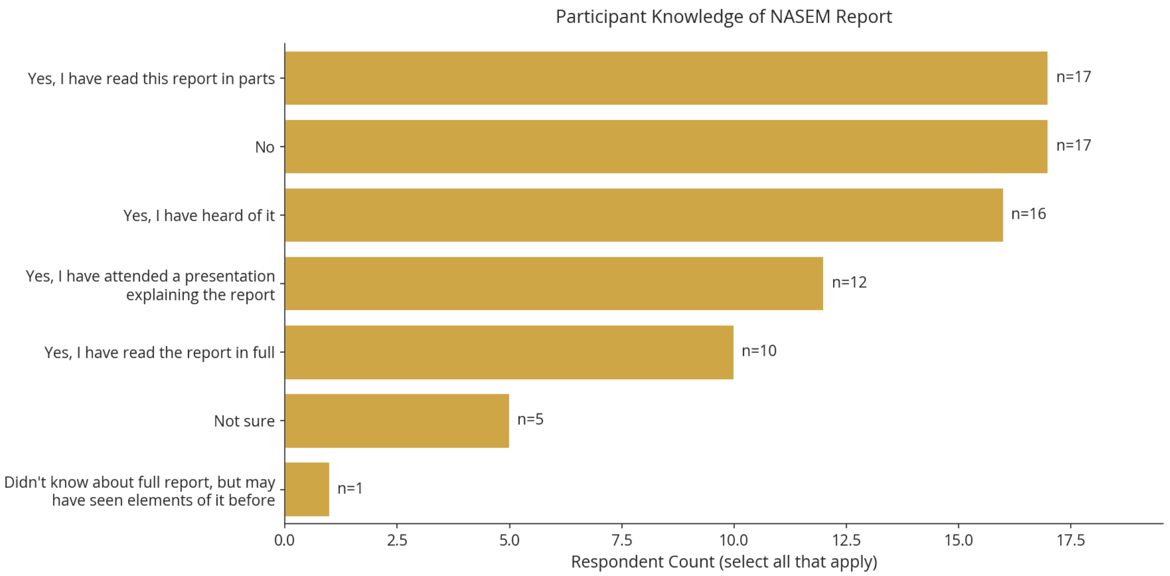


**Supplementary Figure 3:** *Participant self-reported knowledge of the NASEM report.* Participant responses on whether, and to what degree, they were familiar with the NASEM report of population descriptors in human genetics research.

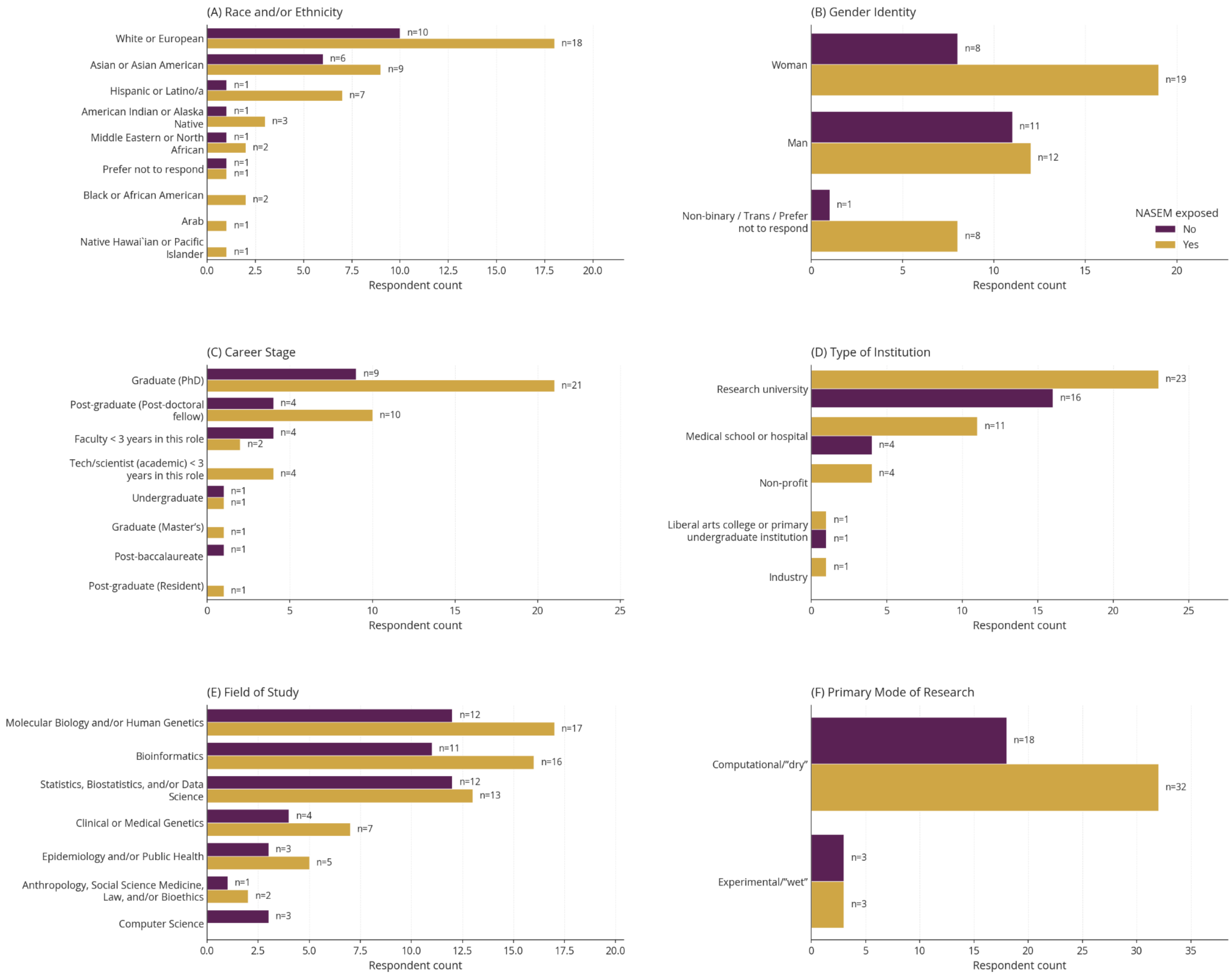


**Supplementary Figure 4:** *Participant demographics by NASEM exposure.* For each participant that chose to respond to demographic questions, the count of those who are NASEM-exposed (n=40) and NASEM-unexposed (n=19) (A) at each career stage, (B) at type of institution, (C) in a specific scientific field, (D) that perform computational or wet lab research; and the participant's (E) self-reported race/ethnicity and (F) gender identity. Participants were allowed to select multiple responses when they felt it was appropriate.

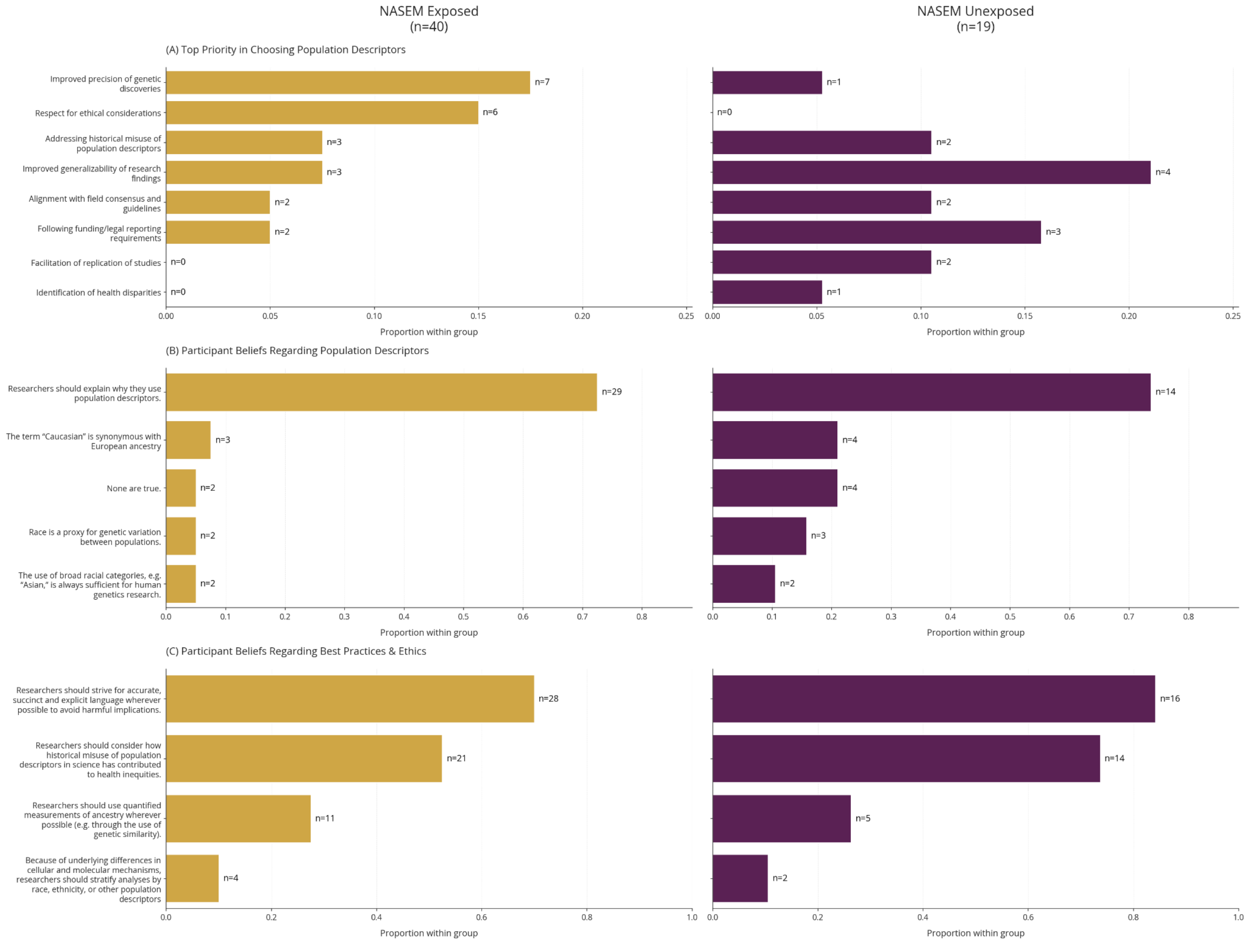


**Supplementary Figure 5:** *Participant beliefs regarding population descriptors.* For the participants that were exposed to the NASEM report (yellow bars) and those unexposed to the report (purple bars), (A) the proportion of participants in that group who ranked each survey option as their top priority, (B) the proportion of participants in each group answering, "*Which of the following do you believe to be true?"*, and (C) the proportion agreeing with statements relating to population descriptor ethics and best practices.

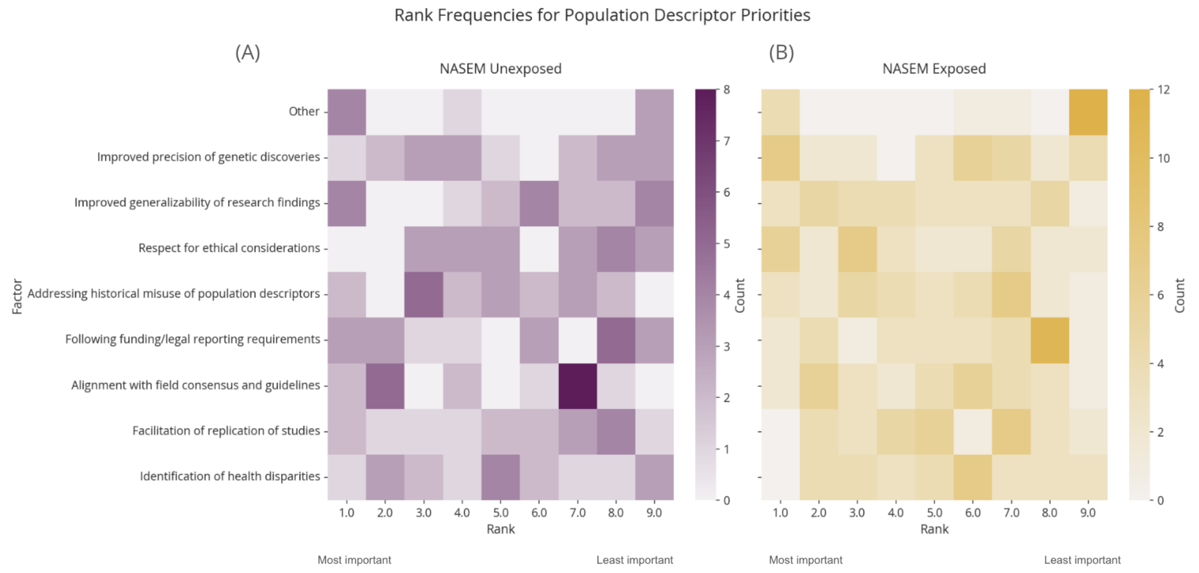


**Supplementary Figure 6:** *Participant priorities in selecting population descriptors.* For the (A) NASEM unexposed and (B) the NASEM exposed, the number of participants selecting 1 (most important) to 9 (least important) for each potential reason to select population descriptors. The darker the color, the greater the number of participants selecting that ranking.

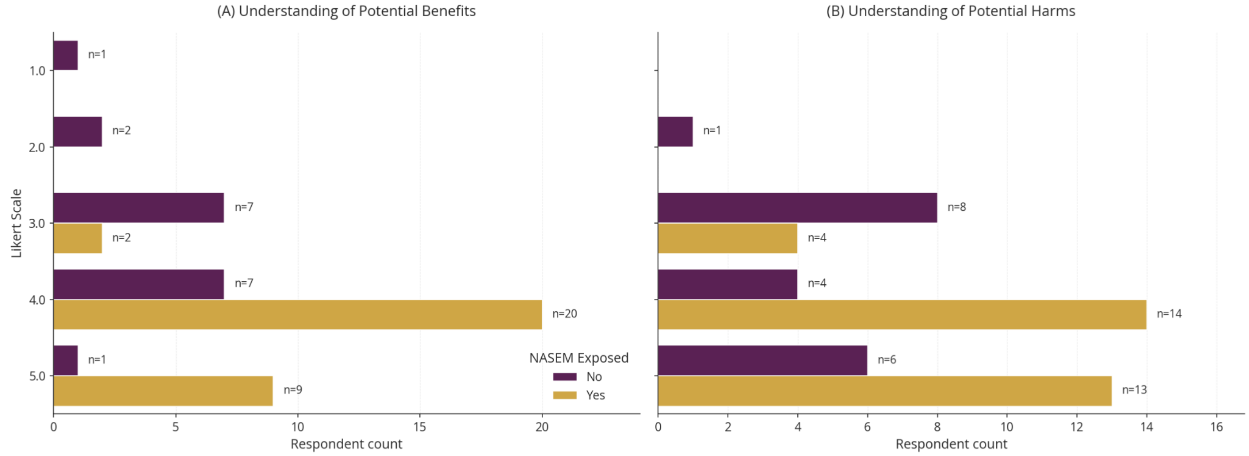


**Supplementary Figure 7:** *Participant confidence in understanding the harms and benefits of population descriptor use.* Participants were asked to rate their understanding from 1-5 (1 being very poor understanding and 5 being very good understanding) of the (A) potential benefits and (B) the potential harms of using population descriptors. Results are disaggregated by NASEM exposure status.

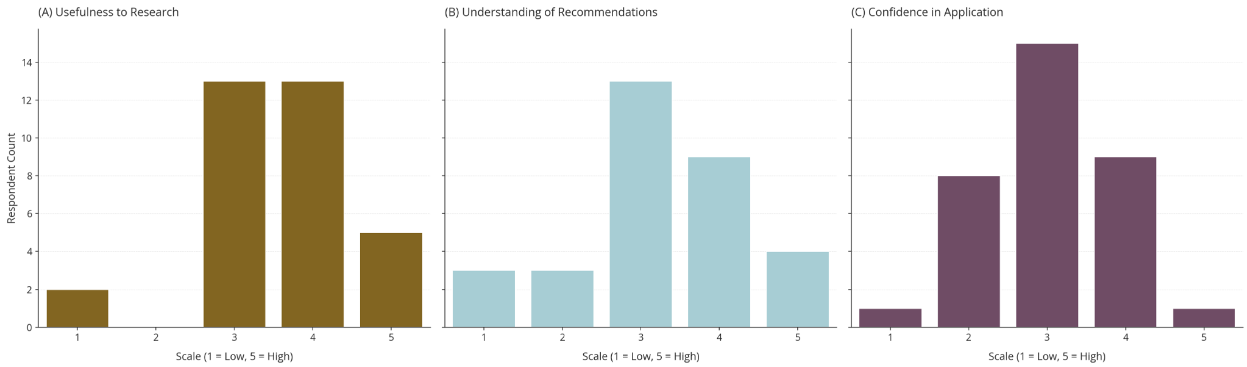


**Supplementary Figure 8:** *Participant attitudes toward the NASEM report.* On a scale of 1 (low) to 5 (high), the number of participants (A) who believe that the NASEM report is useful to their research, (B) who report understanding the NASEM recommendations, and (C) who are confident in their application of the NASEM recommendations in their research.

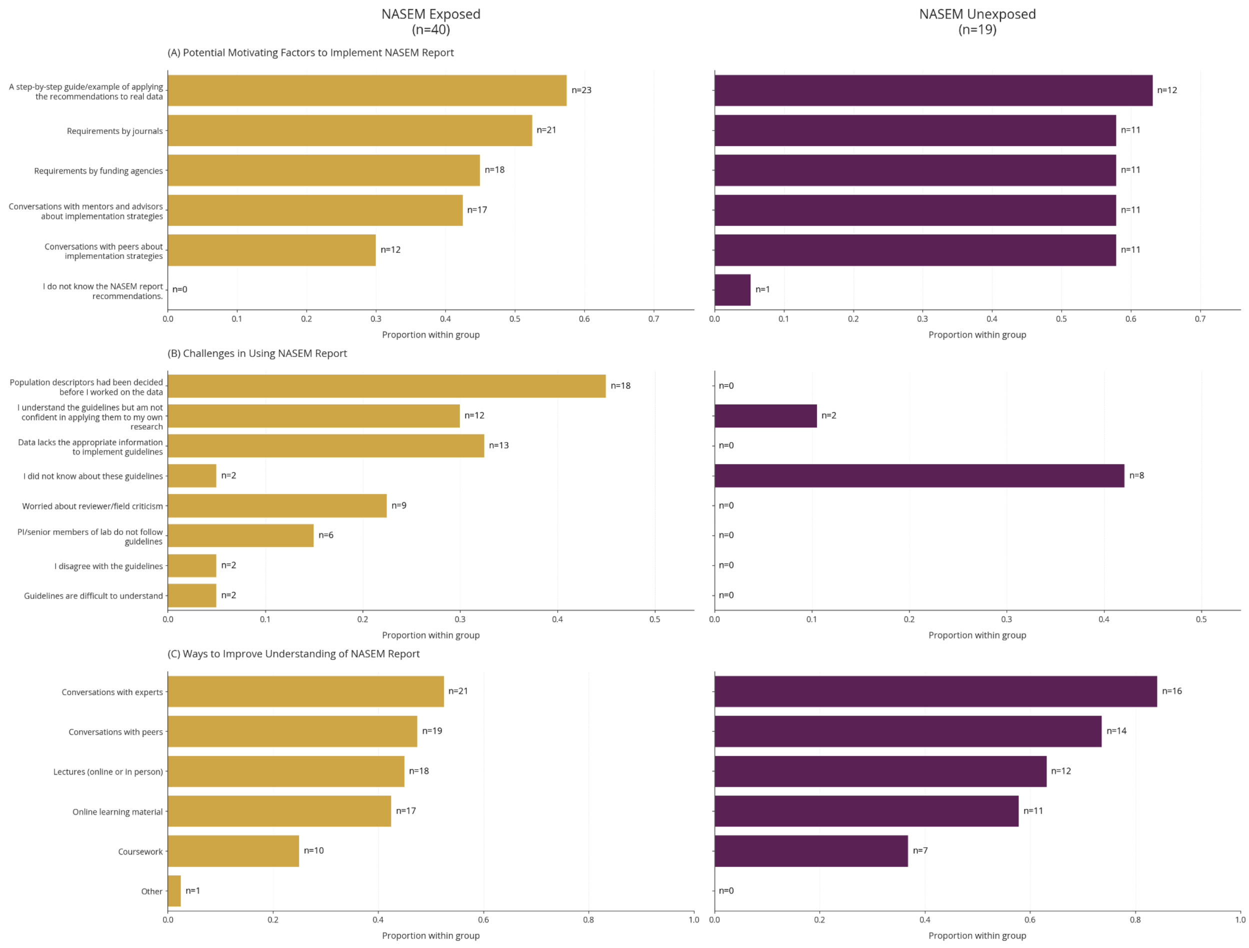


**Supplementary Figure 9:** *Challenges and motivation for using the NASEM report by exposure status.* The proportion of participants with and without NASEM exposure and (A) factors that motivate them to implement the NASEM guidelines in their research, (B) challenges in using the guidelines, and (C) participant suggestions for improving their understanding of the report.

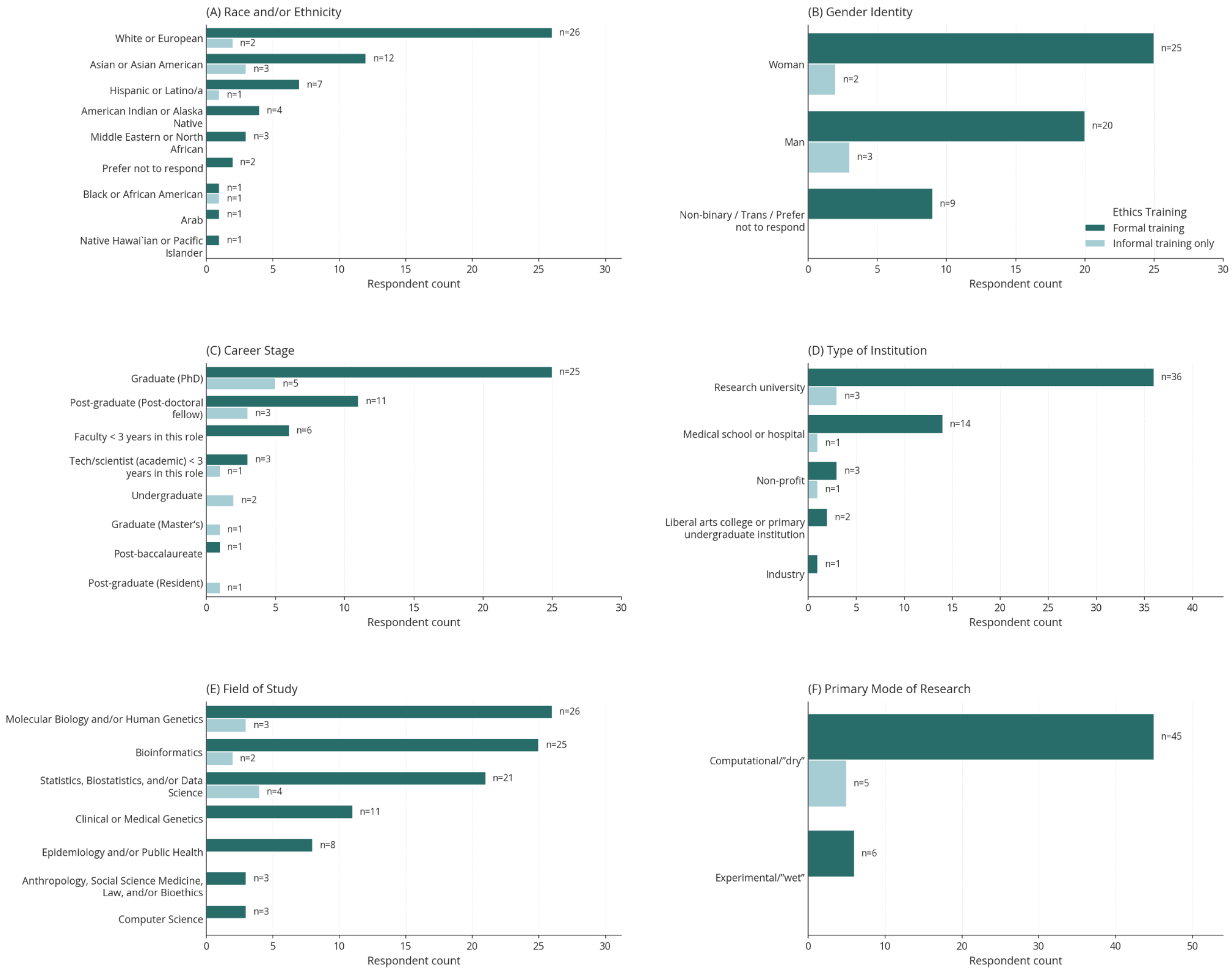


**Supplementary Figure 10:** *Participant demographics by reported ethics training type.* For each participant that chose to respond to demographic questions, the count of those (A) at each career stage, (B) at type of institution, (C) in a specific scientific field, (D) that perform computational or wet lab research; and the participant (E) self-reported race/ethnicity and (F) gender identity. Participants were allowed to select multiple responses when they felt it was appropriate.

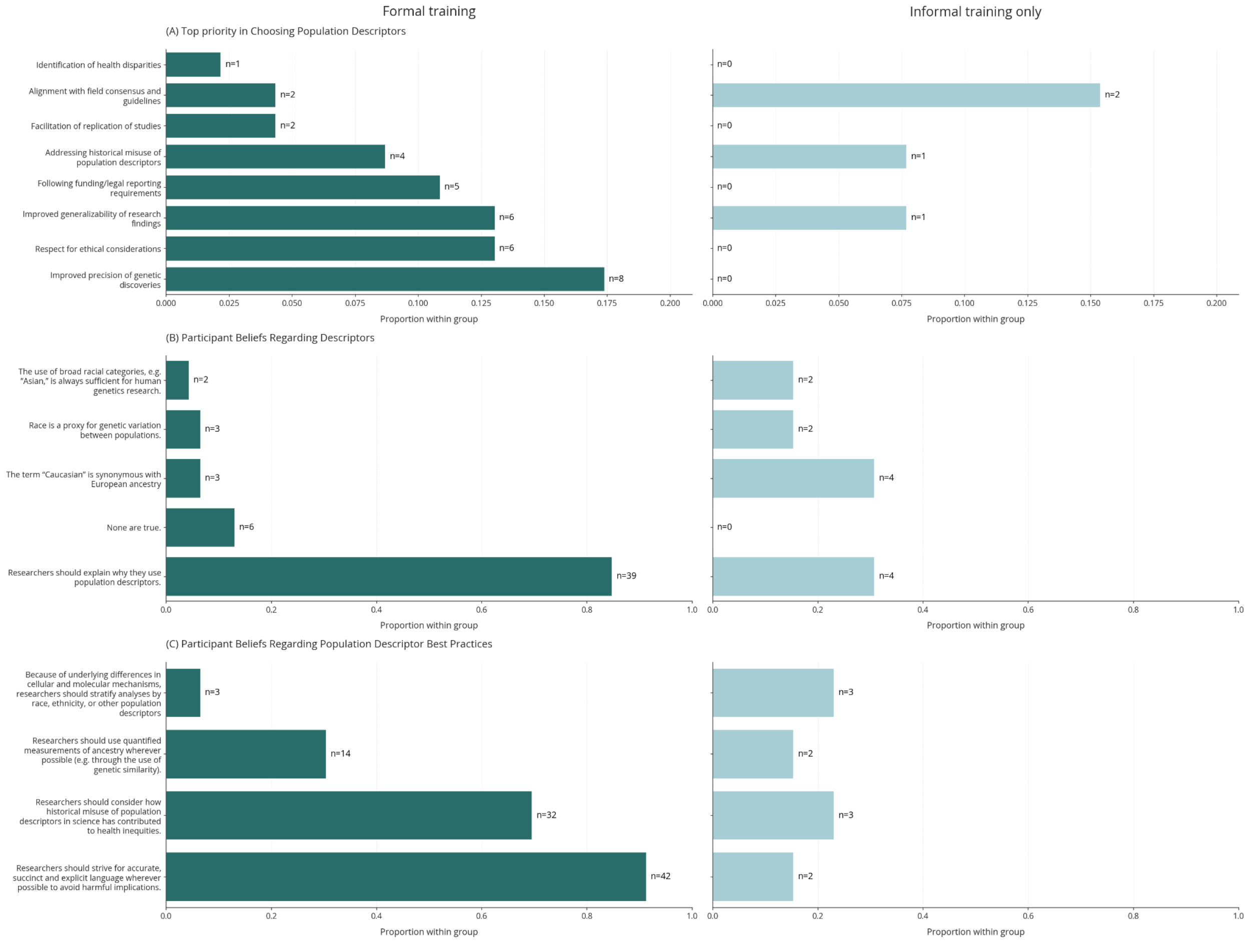


**Supplementary Figure 11:** *Patient beliefs and attitudes toward the NASEM report by ethics training status.* (A) Participant responses to *"Which of the following do you believe to be true?"* (B) Participant beliefs regarding the best practices and ethical considerations when selecting population descriptors. Participants could select multiple options. (C) The top-ranked priority of participants when choosing population descriptors in their research. Participants selected one priority.

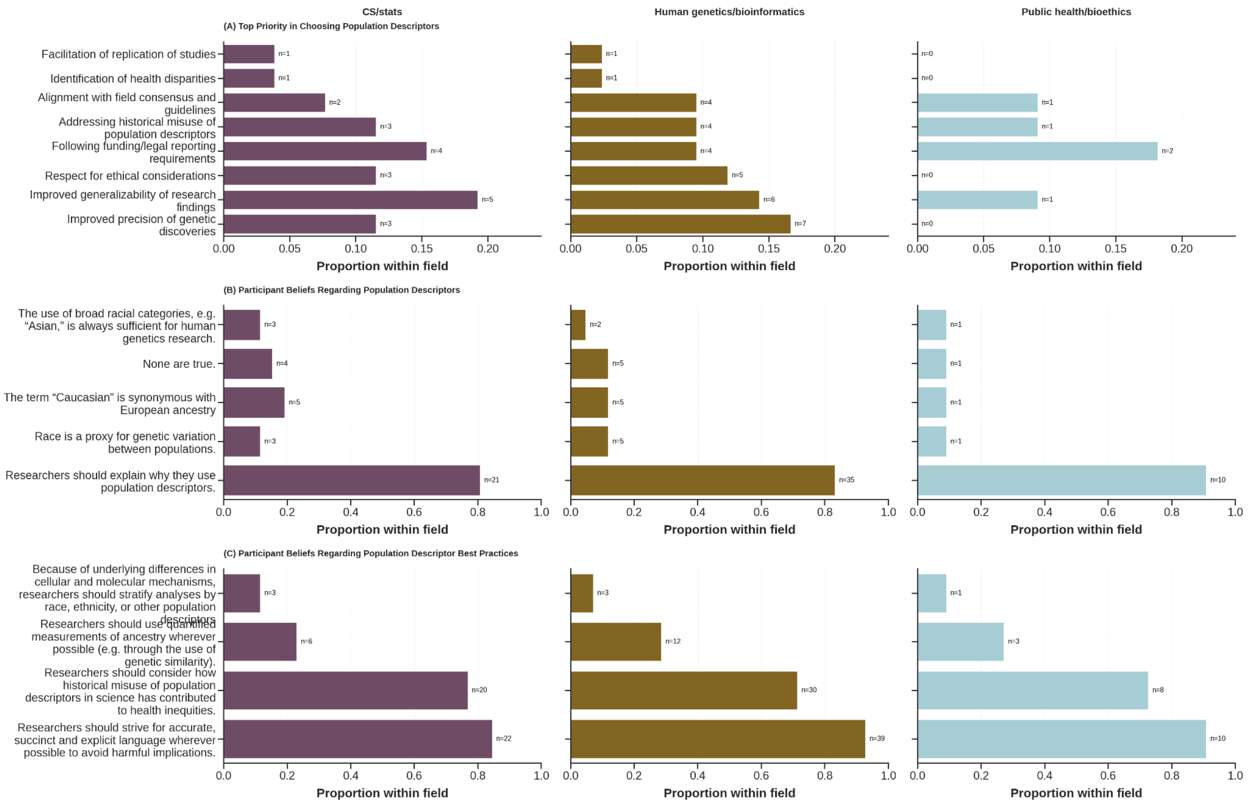


**Supplementary Figure 12:** *Patient beliefs and attitudes toward the NASEM report by training field.* (A) Participant responses to *"Which of the following do you believe to be true?"* (B) Participant beliefs regarding the best practices and ethical considerations when selecting population descriptors. Participants could select multiple options. (C) The top-ranked priority of participants when choosing population descriptors in their research. Participants selected one priority.
